# Targeted Lipid Metabolism Screening Uncovers Regulatory Effects on the STING Immune Response in Mevalonate, Eicosanoid and Fatty Acid Pathways

**DOI:** 10.64898/2026.01.05.697750

**Authors:** Sofia Skobelkina, Ella L Brunsting, Darren J. Perkins

**Author notes:** **Corresponding Author: Darren Perkins**, Howard Hall Room 333. 660 W. Redwood St. Baltimore MD 21201.

## Abstract

The cGAS/STING pathway is a critical signaling hub that orchestrates type I interferon (IFN) responses, autophagy, and programmed cell death in response to double-stranded DNA (dsDNA) or cyclic dinucleotides. While traditionally characterized as a sensor of foreign or mis-localized self dsDNA, recent evidence demonstrates that STING also integrates information about the homeostasis of cellular lipid biosynthesis into the innate inflammatory response. This integration occurs most notably through STING’s sensitivity to *de novo* cholesterol synthesis. However, given that mammalian cells undergo widespread lipid metabolic reprogramming, characterized by alterations in the synthesis of many lipid species in addition to cholesterol, during processes such as malignant transformation to cancer or during infection by intracellular pathogens, we hypothesized that STING function may be regulated by perturbations in other undescribed lipid pathways. To investigate potential other facets of the STING-lipid interface, we have performed a targeted small molecule screen across multiple lipid metabolic pathways, including the mevalonate, PPAR (fatty acid), and arachidonic acid pathways. Our findings reveal that positively and negatively perturbing enzymes within these diverse lipid paths including lipoxygenases and cyclooxygenases can significantly modulate STING-dependent signal transduction and transcriptional programs, identifying metabolic nodes that link lipid homeostasis with innate immune signaling. These results suggest that existing lipid-lowering and metabolic therapies may have unappreciated immunomodulatory effects on STING applicable in cancer and infectious disease, offering new opportunities for therapeutic intervention.

## Introduction

The Cyclic GMP-AMP Synthase/Stimulator of Interferon Genes (cGAS/STING) pathway acts as a central node in immune response and cell biological processes, orchestrating responses related to type I interferon (IFN) signaling, autophagy and programmed cell death^1, 2^. In the resting state, STING is a transmembrane protein on the endoplasmic reticulum (ER) until activated when cGAS binds to double-stranded DNA (dsDNA) and enzymatically synthesizes the cyclic nucleotide second messenger cyclic guanosine monophosphate–adenosine monophosphate (cGAMP). cGAMP binds to a conserved pocket on the cytosolic surface of STING and induces a conformational change triggering STING translocation from the ER to the ER-Golgi intermediate compartment (ERGIC) and the Golgi apparatus through highly regulated vesicle trafficking. STING can also be directly activated independent of cGAS through binding synthetic small molecules^3, 4, 5^ and bacterially derived cyclic second messengers^6^. Once In the Golgi, STING recruits TANK-binding kinase 1 (TBK1), which phosphorylates STING at Serine 365 or 366 (mouse and human respectively). This phosphorylation event licenses the activation of transcription factor IRF3 and leads to a transcriptional program dominated by production of type I interferons^7, 8^.

While most extensively studies as a sensor of dsDNA, emerging research indicates that STING signaling is intricately linked to metabolic health including that of the ER^9, 10^ and mitochondria^11^, and specifically to lipid metabolism related to the synthesis and storage of cholesterol^12^. Sterol Regulatory Element-Binding Protein 2 (SREBP2), the master transcription factor regulating the synthesis of cholesterol through mevalonate pathway^13^, coordinates the strength of STING activation with flux through the mevalonate pathway and reduced cholesterol synthesis has been proposed as a danger signal amplifying the STING dependent IFN response^12^. Mechanistically, pinpointing precisely how ER cholesterol levels are sensed by STING remains ongoing work, however, human disease related perturbations in cholesterol flux including the absence of intracellular cholesterol transporters, such as NPC1, has also been shown to prime STING activity^14^. These observations have led to the hypothesis that STING possesses the capacity to read and integrate “lipid codes” into an innate inflammatory response^12^. Whether other lipid metabolic pathways similarly regulate STING behavior is an open question. However, in addition to cholesterol, other cellular lipid species have also long been known to impact type I interferon induction by immune cells including prostaglandins and leukotrienes, members of the eicosanoid lipid family^15, 16, 17^.

This STING-lipid metabolic interface is particularly relevant in the context of both cancer and infectious disease. In tumor immunology, STING plays a critical role in initiating cell intrinsic anti-tumor immune responses^18, 19, 20^, and therapeutic activation of STING in tumors and the tumor microenvironment is a vital area of drug development and ongoing clinical trials^21^. Within the tumor microenvironment (TME), cGAS/STING activation can act tumor cell intrinsically or through bystander immune cells including dendritic and NK cell activation driving the priming of T cells^22^. Lipid metabolic reprogramming is now widely recognized as a common feature of cancer, allowing malignant cells to adapt to the demands of uncontrolled proliferation and nutrient deprivation^23^. While the shift toward aerobic glycolysis has historically dominated the field of cancer metabolism, recent evidence underscores that alterations in lipid metabolism are equally critical for tumorigenesis^24^. Cancer cells exhibit a distinct “lipogenic phenotype,” characterized by an upregulation of *de novo* fatty acid synthesis, in the presence of abundant exogenous lipids^25^.This shift is often driven by the overexpression of key lipogenic enzymes, such as fatty acid synthase (FASN) and acetyl-CoA carboxylase (ACC)^26^. Concurrently, cancer cells frequently enhance their capacity for fatty acid oxidation (FAO) and lipid droplet storage to maintain cellular homeostasis. Collectively, these perturbations in lipid metabolism support neoplastic growth but may also contribute to local immune suppression^27, 28^. This thinking is supported by pre-clinical and translational data showing that lipid metabolism targeting approaches can have anti-neoplastic effects^29^. Additionally, given that numerous clinically used drugs significantly impact lipid homeostasis and are widely prescribed to treat lifestyle diseases in the western world, how these therapeutics may also be affecting STING-dependent immunity in ways not now understood is a significant question. Similarly, viral and bacterial human pathogens frequently hijack host cell lipid metabolism to support their replication^30^, therefore, the ability of STING to sense lipid metabolic perturbations may also afford greater protection in this context.

To address the gap in understanding of how diverse cellular lipid contexts influence STING, we investigated the impact of small molecule perturbations across several lipid metabolic pathways. In this report, we identify key regulatory effects that link lipid metabolism with immune modulation, suggesting new avenues for therapeutic intervention including mevalonate, PPAR and arachidonic acid pathways.

## Materials and Methods

### Cells

Immortalized Bone Marrow derived macrophages (iBMDM) were a gift of Susan Carpenter (UC Santa Cruz) and have been described previously^31^. iBMDM were maintained in DMEM with 10%FBS and Penn/Strep. Primary Bone Marrow derived macrophages were prepared, as described previously ^32^. Briefly, bone marrow was harvested from C57BL6/J mice and cultured in standard medium supplemented with 20–25% LADMAC cell– conditioned medium ^33^. After 48 hrs, nonadherent cells were removed and adherent cells cultured for an additional 7 days. Adherent cells were harvested using 0.25% Trypsin-EDTA (Life Technologies, Grand Island, NY) and replated for experimental use

### Antibodies and Reagents

Antibodies directed against IRF3 (clone D83B9), P-IRF3 (clone 4D4G), STING (clone D1V5L), P-STING ser 366 (clone D8F4W), P-TBK-1 ser 172 (clone D52C2), p65 (clone D14E12), P-p65 ser 536 (clone 93H1), LC3 A/B (clone D3U4C) and GAPDH (clone 14C10) were purchased from Cell Signaling (Danvers, MA). 3’,2’-Cyclic guanosine monophosphate-adenosine monophosphate sodium salt (cGAMP) was purchased from Tocris (cat no. 7718). Pam3CSK was purchased from Invivogen. Complete mini and PhosSTOP inhibitor cocktail tablets were purchased from Roche (cat no 11836170001 and 4906845001). DMSO, zaragozic acid, methyl-b-cyclodextrin were purchased from Sigma-Millipore. 5-HpETE and LTB4 were purchased from Cayman Chemical and were used following exchange of ETOH for DMSO solvent by evaporation of ETOH under vacuum. Fatostatin A, Ro 48-8071, rosuvastatin, zileuton, ML351, NS398, sc560, GW501516, T0070907, rosiglitazone and 5,6-Dimethyl-9-oxo-9*H*-xanthene-4-acetic acid (DMXAA) were purchased from Tocris.

### Lipid Pathway Screening

For the small molecule screen of lipid pathways, immortalized BMDMs were plated in 12 well tissue culture treated plates at a density of 3 x10^5^/well. After adherence overnight, iBMDMs were pretreated with individual compounds dissolved in DMSO for 3 hours at a concentration of 10μM. Following pretreatment individual wells were stimulated with DMXAA at 10μg/ml for an additional 3 hours. Cells were then harvested and processed to determine gene expression using qRT PCR method.

### Viability Assay

To determine impacts of small molecules on cell viability, iBMDMs were treated with individual compounds at 10μM in quadruplicate for the times indicated in the screening assay and viability was assessed using the CellTiter-Glo kit (Promega) according to manufacturers instructions.

### Quantitation of secreted cytokines

Cytokine/chemokine levels in macrophage culture supernatants were analyzed by enzyme linked immunosorbent assay (ELISA) at the Cytokine Core Laboratory (University of Maryland, School of Medicine).

### qRT PCR

Total mRNA was isolated from cells using TRIPure (Sigma/Roche Cat No. 11667165001) reagent, according to manufacturer’s instructions. A total of 1 μg RNA was used in cDNA synthesis using the iScript cDNA synthesis kit (BioRad) according to manufacturer’s instructions. qRT-PCR was performed on cDNA using an Applied Biosystems Quant Studio 3 system with POWER SYBR Green reagent (Applied Biosystems) and transcript-specific primers as previously described^34^. Levels of mRNA for specific genes are reported as relative gene expression normalized to that of untreated cells (“fold induction”;^35^). The housekeeping gene encoding Glyceraldehyde 3 phosphate dehydrogenase (GAPDH) was used for normalization of RNA levels within each sample. In addition to previously published primer sequences for detection of cytokine gene expression, the following primer sets were used to detect mRNA: (GAPDH FWD-AGC CTC GTC CCG TAG ACA AAAT; REV-TGG CAA CAA TCT CCA CTT TGC) (IFN-β FWD-CAC TTG AAG AGC TAT TAC TGG AGG G; REV-CTC GGA CCA CCA TCC AGG) (IP10 FWD-CCA CGT GTT GAG ATC ATT GCC; REV-GCC CTT TTA GAC CTT TTT TGG C) (TNF FWD-GACCCTCACACTCAGATCATCTTCT; REV-CCACTTGGTGGTTTGCTACGA) (IL6 FWD-TGTCTATACCACTTCACAAGTCGGAG; REV-GCACAACTCTTTTCTCATTTCCAC)

### Quantitation of secreted cytokines

Cytokine/chemokine levels in macrophage culture supernatants were analyzed by enzyme linked immunosorbent assay (ELISA) at the Cytokine Core Laboratory (University of Maryland, School of Medicine).

### Immunoblot analysis

Immunoblotting was carried out substantially as previously described^36^. Whole-cell lysates from macrophage cultures were obtained by the addition of BI lysis buffer (20 mM HEPES, pH 6.8, 1.0% Triton X-100, 0.1% SDS, 150 mM NaCl, and Complete mini and PhosSTOP tablets) and subsequent incubation at 4°C. Cell lysates were separated by denaturing SDS-PAGE and subsequent transfer to polyvinylidene fluoride (PVDF) membranes. Blots were incubated overnight in relevant primary antibodies (identified above) at 4°C, were washed three times with phosphate buffered saline (PBS), and then were incubated with the appropriate horseradish peroxidase (HRP)-conjugated secondary antibody [Jackson HRP anti-rabbit (catalog number 111-035-003) or Jackson HRP anti-mouse (catalog number 115-035-003i), Jackson Immuno-chemicals]. Blots were developed following incubation in the ECL Plus Western Blotting Detection Reagent (Amersham Bioscience).

### Statistical analysis

Statistical analysis, when applied, was done using the Prism software v5.0. All *t*-tests were done using a two-tailed analysis. A p-value <0.05 was considered statistically significant.

## Results

### Perturbations in Multiple Lipid Pathways Affect STING Induction of IFN-β

To identify unknown impacts of lipid metabolic pathways with previously described immune relevance on STING receptor functionality we carried out a targeted small molecule screen utilizing immortalized mouse C57BL6/J bone marrow derived macrophages (iBMDMs). We focused our initial screen on the well described and clinically relevant mevalonate (cholesterol synthesis), arachidonic acid (cyclooxygenase and lipoxygenase), and peroxisome proliferator activated receptor (PPAR) pathways as each has significant qualitative and quantitative impacts on macrophage lipid metabolism and are known to be differentially engaged during macrophage immune polarization. iBMDMs were pretreated for three hours in triplicate wells in culture either with vehicle alone (DMSO) or methyl beta cyclodextrin, a molecular lipid sink with preference for cholesterol binding, as controls, or with individual compounds acting as antagonists or agonists of enzymes in each of the three metabolic pathways. Following three hours of pre-treatment, iBMDMs were subsequently stimulated with the synthetic mouse STING agonist 5,6-Dimethylxanthenone-4-acetic Acid (DMXAA 10μg/ml) for an additional three hours to drive STING transcriptional responses. Following STING activation, cells were analyzed for the induction of type I interferon (IFN-β) by qRT-PCR. Quantitation of IFN-β message revealed that compounds affecting the activity of enzymes in each of the three lipid regulatory pathways were capable of enhancing STING dependent IFN induction, effects which have not been previously described for the majority of these molecules (Figure 1A). The compound which consistently displayed the largest enhancement of STING responses was Fatostatin A, a pre-clinical drug with demonstrated impacts in models of dyslipidemia and prostate cancer^37, 38^. Fatostatin is an antagonist of the transcription factor SREBP2, which is the master regulator of expression for enzymes in the mevalonate pathway which governs cholesterol production^13^. Positive effects of Fatostatin on STING responses in our screen are congruent with published results showing enhanced STING responses in genetic models of reduced cholesterol synthesis^12^. Importantly, two clinically approved compounds: zileuton (brand name zyflo) an antagonist of 5-lipoxygenase (5-LO) prescribed to treat asthma in patients^39^, and rosiglitazone (brand name Avandia) a PPARγ agonist used clinically to treat type 2 diabetes^40^ both significantly altered STING dependent IFN induction. Neither zileuton nor rosiglitazone have been previously reported to regulate the cGAS/STING pathway. While zileuton increased STING responses, somewhat surprisingly, rosiglitazone was the only compound in our screen identified as a suppressor of STING (Fig 1A). Enzymatic targets for each compound and the relative direction of effects on STING response are indicated in Figure 1B. The effect of each compound on iBMDM cell viability over the relevant time period were assessed and found to be negligible (Figure 1C). Measurement of secreted IFN-β protein levels in culture media were consistent with conclusions from mRNA quantitation (Figure 1D). While the iBMDM cell line used in our initial screen is not malignantly transformed and has growth characteristics similar to primary cells, it is formally possible that the process of immortalization altered relevant metabolic processes so we recapitulated select drug responses in primary C57BL6/J BMDMs and found them consistent with iBMDM results (Figure 1E). DMXAA is a synthetic small molecule which is directly cell permeable and a strong STING agonist compared to endogenous activators^4, 41^. To determine whether physiological STING activators could also be affected by lipid metabolism modifying compounds, iBMDMs were pre-treated with Fatostatin A followed by stimulation with the endogenous STING ligand cyclic guanosine monophosphate-adenosine monophosphate (cGAMP 10μg/ml) for three hours. Fatostatin A significantly increased both IFN-β and IP10 transcription in response to cGAMP (Figure 2A). The mechanism by which alterations in cholesterol flux prime STING activity is still being worked out and multiple mechanisms have been proposed^12, 42, 43^. To determine whether pharmacologic treatment with Fatostatin A produced a generalizable enhancement in transcriptional responses driven by innate pattern recognition receptors or was specific to STING, iBMDMs were pretreated with Fatostatin A and subsequently stimulated with the TLR2 agonist PAM3CSK4 or recombinant tumor necrosis factor (TNF). Measurement of induced IL-6 and TNF transcription indicated that neither TLR2 nor TNFR responses were similarly primed by Fatostatin A (Figure 2 B and C). Transcription of STING dependent immune response genes such as IFN-β and IP10 is governed by activity of multiple common signaling pathways and transcription factors (e.g. NF-κB and IRF3) in addition to regulated receptor post translational modifications. STING additionally induces a cell intrinsic autophagic/xenophagic response independent of its downstream cytokine transcriptional program and whether the compounds we identified in our screen were acting at the level of the STING receptor itself or were preferentially modulating a single signaling node downstream of STING is a relevant consideration addressing mechanism. To determine effects of compounds we identified which both positively and negatively impact STING transcription on primary immune response signaling pathways, we performed western blot characterization during a time-course following DMXAA stimulation of iBMDMs pretreated with vehicle (DMSO), Fatostatin A, zileuton or rosiglitazone. In agreement with our mRNA expression data, both Fatostatin A and zileuton generated an increase in STING receptor activation as assayed by phosphorylation of serine 365 arguing that these compounds enhance responses at the receptor level. Congruent with this, we observed moderate increases above vehicle treatment in both IRF3 and NFκB p65 transcription factor phosphorylation in response to DMXAA (Figure 3 A & B). The PPARγ agonist rosiglitazone, in contrast, produced marked suppression of both STING receptor phosphorylation and downstream signaling pathways (Figure 3C). Of note, the effects of rosiglitazone on the STING induced autophagic response, as assayed by the appearance of the lipidated form of the LC3 protein (LC3II), were profoundly suppressive, arguing that rosiglitazone antagonizes both transcriptional responses and cell intrinsic host defense pathways (Figure 3C). Zileuton is a selective antagonist of the 5-Lipoxygenase enzyme which acts on arachidonic acid liberated from the phospholipid bilayer in response to inflammatory stimuli to produce secondary lipid species which can then be further metabolized to generate the secreted immune-active lipid leukotrienes^44^. The proximal product of 5-LO action on arachidonic acid is the poly unsaturated fatty acid 5-HpETE (see Figure 4A). As lipoxygenase pathways have not been explicitly linked to STING regulation previously, it was of interest to confirm the results of our inhibitor study and to begin to address the mechanism of 5-LO interaction with STING by determining whether 5-HpETE itself was sufficient to inhibit STING. Pre-loading of iBMDMs with 5-HpETE prior to stimulation with DMXAA led to reduced IFN-β and IP10 transcription (Figure 4B). 5-LO generated 5-HpETE can be subsequently metabolized into many secondary lipids with the most well characterized 5-LO product being the secreted immunomodulatory leukotrienes which have manifold effects on inflammatory responses^45^. LTB4 is the predominating leukotriene specie produced by mouse macrophages in a number of models^46^ and so we asked whether LTB4 feedback may be ultimately responsible for STING regulation by 5-LO. Pre-treatment of iBMDMs with escalating doses of LTB4 did not however produce a modulation of IFN-β transcription in response to DMXAA indicating that the relevant lipid specie downstream of 5-HpETE remains to be identified (Figure 4C). The relevant lipid pathways and the target enzymes for the compounds in this study are described in (Figure 4D).

**Figure 1.**
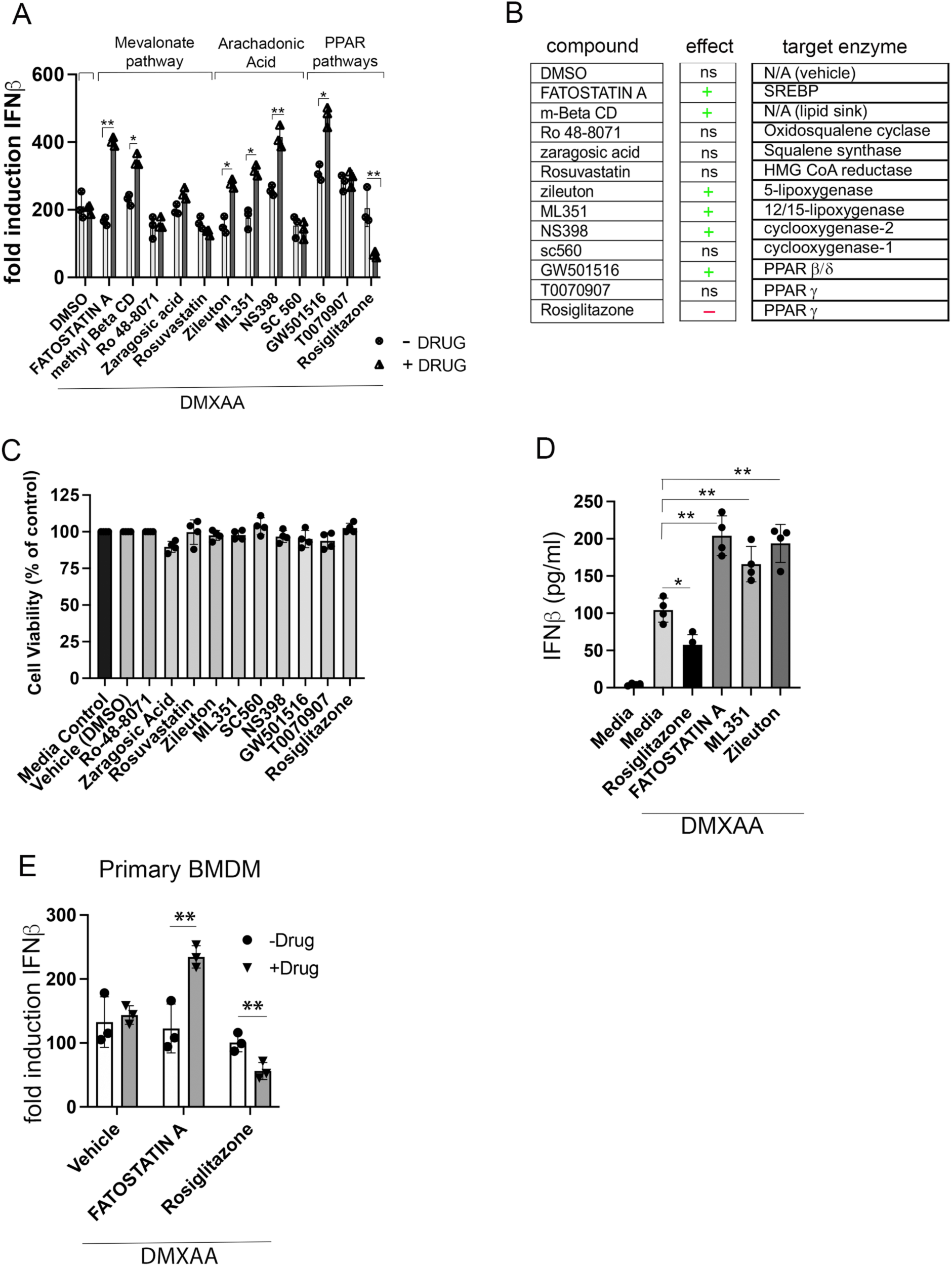
Perturbations in Multiple Lipid Metabolic Pathways Affect STING Induction of IFN-β. A) iBMDMs pretreated for 3 hours with the indicated compounds were stimulated with the synthetic STING agonist DMXAA for a subsequent 3 hours and total RNA was harvested for quantification of IFN-β message by qRT-PCR *=p≤.05, **p<.01. B) Summary of the target enzymes and effects on STING of compounds used in (A). C) iBMDMs treated with vehicle (DMSO) or the individual compounds used in (A) were assayed for viability using the CellTiter-Glo kit. D) Cell media harvested from iBMDM wells treated with individual compounds from (A) were assayed for levels of IFN-β protein by ELISA. E). Primary BMDMs generated from 6-8 week C57BL6/J mice were treated with indicated compounds for 3 hours followed by DMXAA for a subsequent 3 hours and total RNA was harvested for quantification of IFN-β message by qRT-PCR. N > 3.

**Figure 2.**
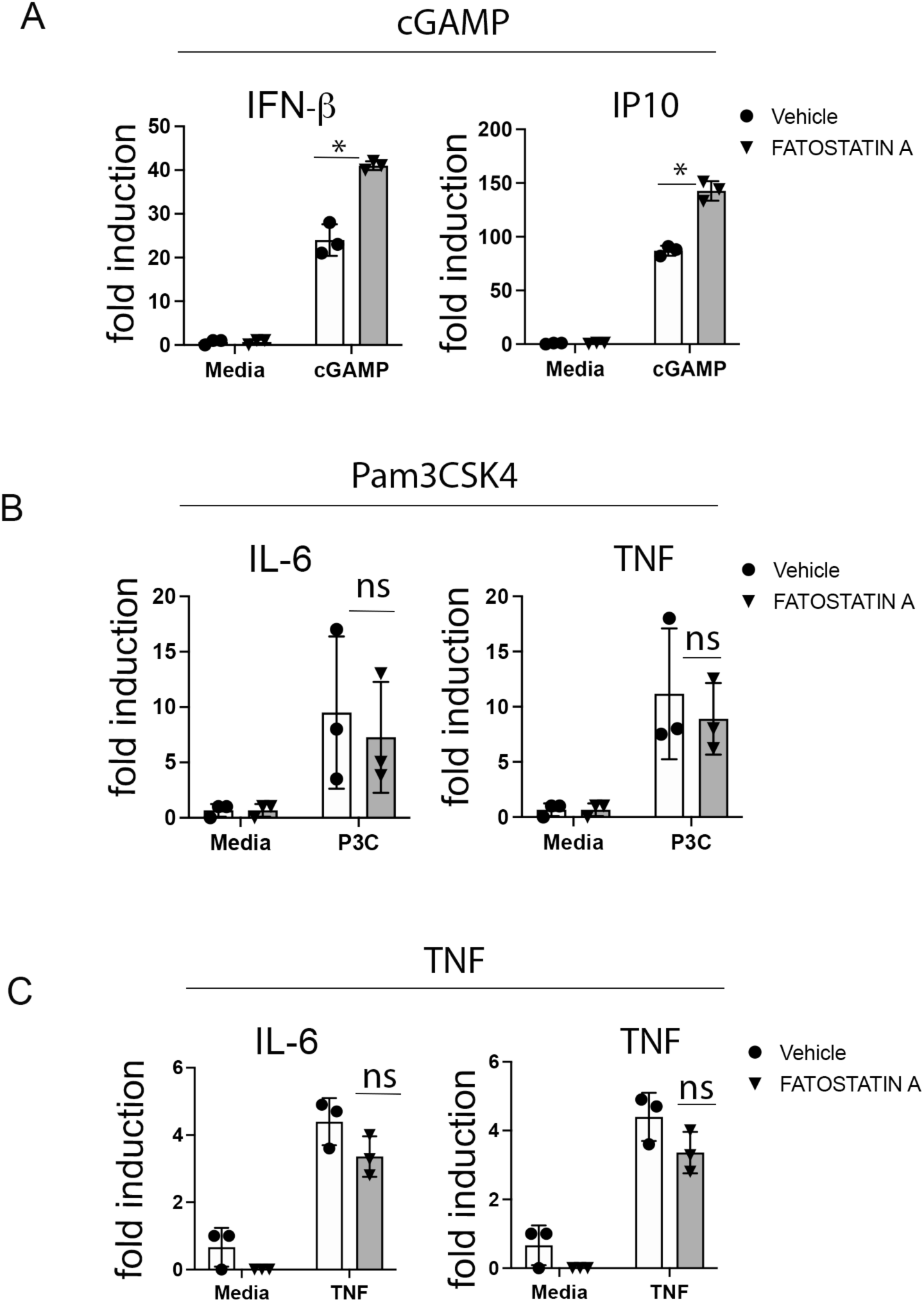
Inhibiting Mevalonate Pathway flux through SREBP Antagonism Preferentially Affects STING PRR Transcriptional Responses. A) iBMDMs were pretreated for three hours either with vehicle (DMSO) or FATOSTATIN A (10µM) followed by stimulation with cGAMP (10µg/ml) added to culture media for an additional 3 hours. Total RNA was harvested for quantification of IFN-β and IP10 mRNA by qRT-PCR. B) iBMDMs were pretreated for three hours either with vehicle (DMSO) or FATOSTATIN A (10µM) followed by stimulation with synthetic TLR2/1 agonist Pam3CSK4 (100 ng/ml) added to culture media for an additional 3 hours. Total RNA was harvested for quantification of IL-6 and TNF mRNA by qRT-PCR. C) iBMDMs were pretreated for three hours either with vehicle (DMSO) or FATOSTATIN A (10µM) followed by stimulation with recombinant TNF (100 ng/ml) added to culture media for an additional 3 hours. Total RNA was harvested for quantification of IL-6 and TNF mRNA by qRT-PCR. N = 3. *p<.05.

**Figure 3.**
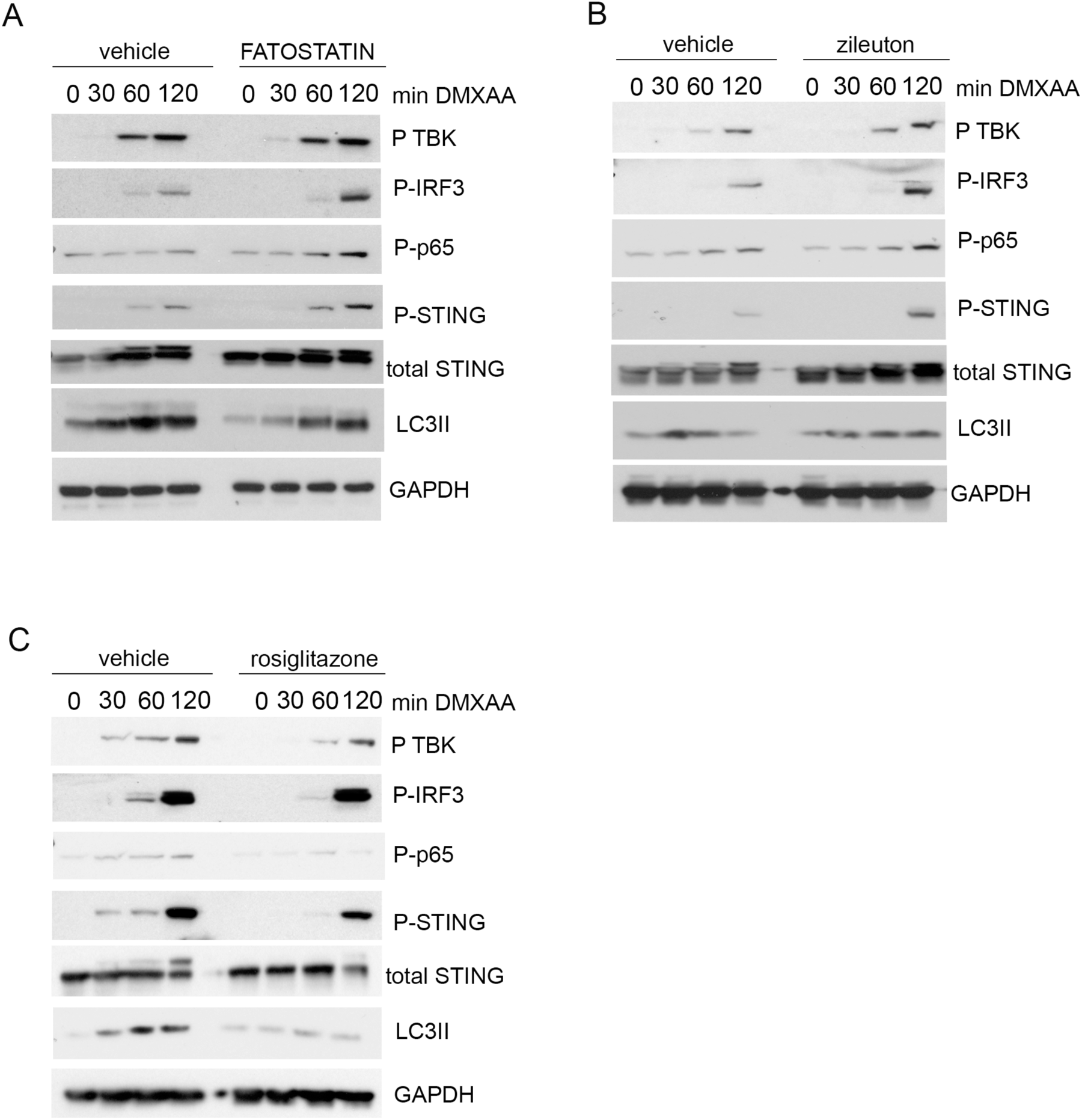
Small Molecule Perturbation of Mevalonate, 5-LO and PPARγ Lipid Pathways Broadly Affects STING Receptor Activation and Signaling. A-C) iBMDMs were pretreated for three hours either with vehicle (DMSO), FATOSTATIN A (10µM) (A), zileuton (B) (10µM) or rosiglitazone (C) (10µM) followed by stimulation with DMXAA (10µg/ml) for the indicated times. Whole cell lysates were harvested and analyzed by immunoblot using antibodies directed against the indicated protein targets. Representative of n= 5.

**Figure 4.**
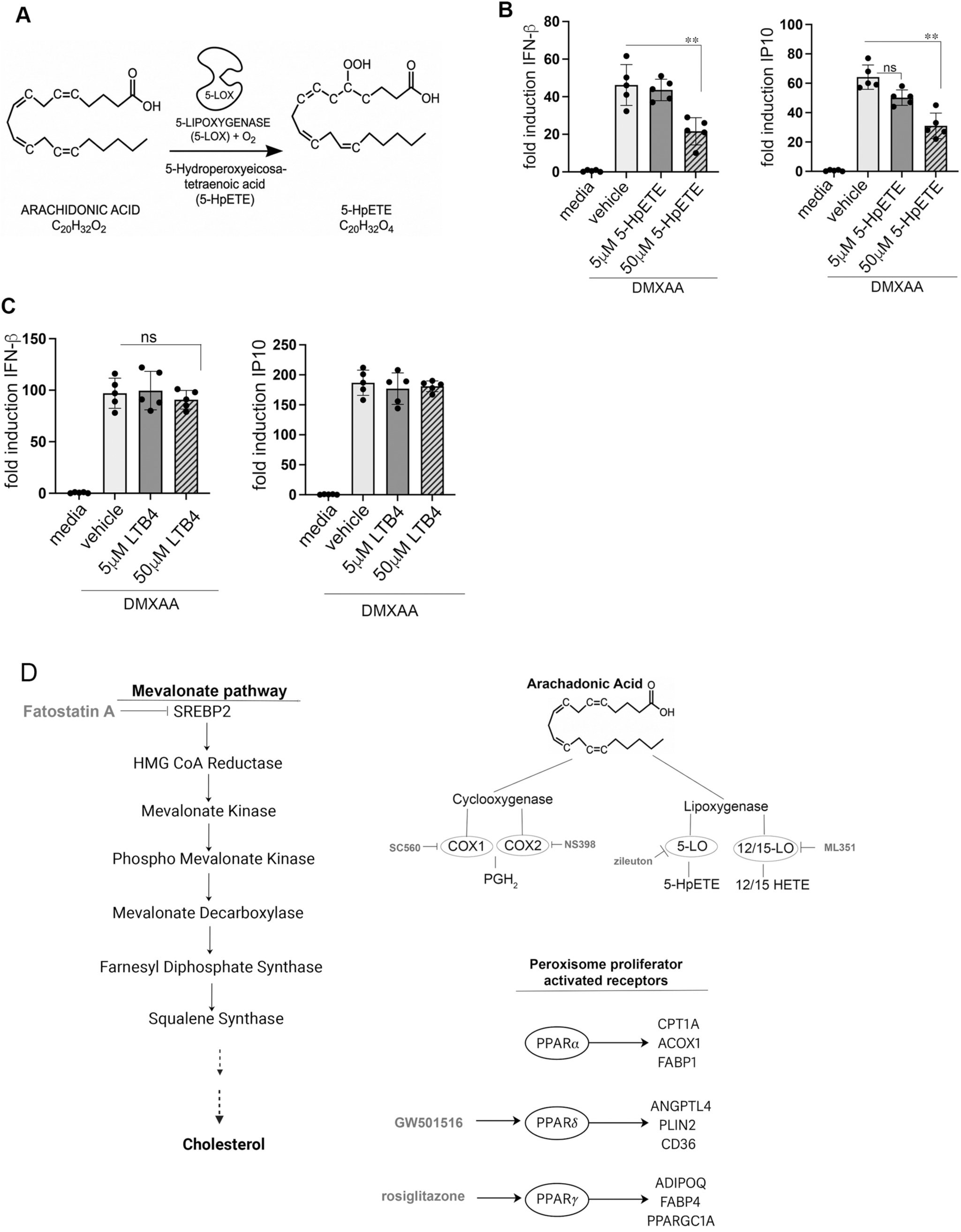
5-LO lipid pathway metabolite 5-HpETE but not Leukotriene B4 Suppresses STING Transcriptional Responses. A) Model of 5 Lipoxygenase enzymatic action on arachidonic acid to produce the proximal lipid metabolite 5-hydroperoxyeicoasatetranoic acid. B) iBMDMs were pretreated for three hours either with vehicle (DMSO), 5µM 5-HpETE or 50µM 5-HpETE followed by stimulation with DMXAA (10µg/ml) added to culture media for an additional 3 hours. Total RNA was harvested for quantification of IFN-β and IP10 mRNA by qRT-PCR. **p<.01 C) iBMDMs were pretreated for three hours either with vehicle (DMSO), 5µM LTB4 or 50µM LTB4 followed by stimulation with DMXAA (10µg/ml) added to culture media for an additional 3 hours. Total RNA was harvested for quantification of IFN-β and IP10 mRNA by qRT-PCR. D) Schematic models of mevalonate, arachadonic acid, and peroxisome proliferator activated receptor pathways indicating locations of relevant targets for compounds used in this study.

## Discussion

In this study we have undertaken a targeted small molecule based functional screen in immortalized and primary mouse macrophages to test the hypothesis that the strength of STING activation to a given dose of ligand is interconnected with the state of multiple immune regulating lipid pathways. Our findings expand upon the evolving characterization of STING as a multi-modal metabolic sensor, moving beyond its well-established role as a direct cyclic dinucleotide receptor. While previous studies, primarily utilizing genetic models, have identified such cell biological contexts impacting STING signaling, our report identifies additional druggable nodes that leverage the intersection of lipid metabolism and STING dependent innate immunity.

Our data builds on the landmark work from York et al.^12^, and subsequently others, proving that STING receptor function is inextricably linked to broader lipid metabolism, most conspicuously *de novo* cholesterol synthesis through the mevalonate pathway. Although the precise molecular conduits connecting mevalonate pathway intermediates to the STING scaffold remain to be fully elucidated, our data further reinforces this biological connection. We identify in this current study the small molecule Fatostatin A, an antagonist of SREBP2, as a compound which can be used to leverage changes in the mevalonate pathway to enhance STING downstream signals and transcriptional responses. Previous literature has noted anti-tumor efficacy of Fatostatin A, and related clinically utilized statins, across various malignancies, most prominently prostate cancer, though a single underlying mechanism for this effect has remained elusive. Our results suggest that the potentiation of the cGAS-STING axis could represent an additional underappreciated mechanism of action for these compounds, providing a pro-inflammatory stimulus within the tumor microenvironment (TME) that could bypass tumor-mediated immune evasion. We additionally observed that two FDA approved and clinically prescribed drugs which target distinct lipid paths related to immune metabolism, zileuton and rosiglitazone, could amplify or suppress STING responses respectively. This data provides an initial description of two additional host directed pharmacological tools for manipulating STING in disease models. Our data also, suggests that STING modulation could, at least in part, underlie the beneficial responses observed with these drugs when used to treat asthma and diabetes respectively.

From a more fundamental innate immunology perspective, York et al. hypothesized that STING reads perturbations in mevalonate pathway flux as a lipid code and integrates that coded information into the innate inflammatory response. Whether and what other lipid codes may exist and how they might mechanistically communicate with cGAS/STING remains an ascendent area of work in the field. A significant finding of our screen is that inhibiting any of the immune associated enzymes involved in metabolizing arachidonic acid (AA) (5-LO, 15-LO, and COX-2) primes STING activation and subsequent type I Interferon (IFN) production. This is particularly notable given that STING physically associates with and regulates FADS1/2, rate-limiting enzymes in arachidonic acid biogenesis^47^. These data may suggest that the level of the cellular pool of AA itself is an additional lipid code to which STING is sensitive.

Having such an AA lipid code could be useful to mammals given STING’s role in natural innate cancer immune-surveillance, as the rate of AA turnover is accelerated by orders of magnitude in certain tumor types^48^. We specifically found that zileuton, an antagonist of 5-LO, the enzyme which acts on AA to produce the poly unsaturated fatty acid 5-HpETE, potentiates STING activity. This is intriguing in light of the fact that 5-LO expression and activity are vastly overexpressed in some cancer types^49^. Having a STING regulatory circuit tied to the levels of AA may also be a benefit as some of the immunomodulatory lipids produced as a result of AA metabolism are hyper produced by cancer cells and contribute to immunosuppressive tumor microenvironments. In addition to the AA associated compounds, we identified the transcription factor PPARγ agonist rosiglitazone as a suppressor of STING activation. PPARγ is known to regulate many intersecting lipid pathways but can also increase cholesterol synthesis and storage. Rosiglitazone driving PPARγ activity may therefore feed forward into the already described STING-cholesterol axis. While STING is activation is clearly sensitive to the upstream lipid metabolic context, it should be noted that STING activation drives a potent downstream production of type I interferon which can further rewire lipid metabolism^50^. Interferon can also display broad immunosuppressive properties further exacerbating the impacts of changes to lipids biosynthesis^34, 51^.

The idea of STING regulating lipid codes is congruent with the emerging conception of homeostasis altering molecular processes (HAMPS) as innate immune response generating danger signals. While the classic models of immunology focused on pathogen and damage generated signals (PAMPs and DAMPs), HAMPs describe the immune system’s ability to monitor the functional state of a cell rather than merely the presence of molecular structures. HAMPs are therefore defined as perturbations in cellular or physiological homeostasis that that signal a system is no longer operating within its “set point”^52^. The intersection of HAMPs and metabolism (immunometabolism) is an active area of current research, and several recent examples of this interconnectivity have come to light. For example disruptions in the mevalonate pathway are sensed as a HAMP by the inflammasome system as inhibition of HMG-CoA reductase (HMGCR) by pathogens or drugs leads to a drop in downstream isoprenoids, which is sensed by NLRP3 and initiates inflammasome priming^53^. Secondly, recent studies have identified the lipid sphingosine-1-phosphate (S1P) as a metabolic HAMP. Alterations in S1P levels, caused by cellular stress or infection, are sensed by NOD1/2 receptors. This identifies NOD receptors not just as peptidoglycan sensors, but as metabolic hubs that monitor lipid homeostasis^54^.

In addition to malignant diseases, lipid metabolism targeting compounds affecting STING may also be useful in developing new tools to treat inflammation associated lifestyle diseases. Development of synthetic STING agonists/antagonists to treat lifestyle associated diseases, most notably Metabolic Dysfunction Associated Steatohepatitis (MASH) is an active next generation frontier in pharmaceutical drug development^55, 56^. In MASH, systemic dyslipidemia leads to lipid loading of liver cells which drives mitochondrial dysfunction and ultimately mitochondrial DNA leakage producing a chronic low grade cGAS/STING activation signature^57^.

With a new focus on STING moving away from a goal of broad inhibition/activation toward a selective targeting that preserves STING’s beneficial housekeeping functions while quenching chronic, sterile inflammation. Having an armamentarium of host directed small molecules which may differentially tune STING function in response to a subsequent stimulation without entirely turning STING ‘On’ or ‘Off’ may be an advantageous direction for the future.

## Funding

The authors declare this work was supported by the National Institute of Allergy and Infectious Disease, National Institutes of Health (Grants R21AI152051 and R21AI180181 to D.J.P.) Additional support was provided by funds through the Maryland Department of Health’s Cigarette Restitution Fund Program CH-649-CRF. ELB was supported by T32AI095190.

## Notes

### Competing Interest Statement

The authors have declared no competing interest.

## References

1. Motwani, M., Pesiridis, S. & Fitzgerald, K.A. DNA sensing by the cGAS-STING pathway in health and disease. Nat Rev Genet 20, 657–674 (2019).

2. Hopfner, K.P. & Hornung, V. Molecular mechanisms and cellular functions of cGAS-STING signalling. Nat Rev Mol Cell Biol 21, 501–521 (2020).

3. Roberts, Z.J. et al. The chemotherapeutic agent DMXAA potently and specifically activates the TBK1-IRF-3 signaling axis. J Exp Med 204, 1559–1569 (2007).

4. Prantner, D. et al. 5,6-Dimethylxanthenone-4-acetic acid (DMXAA) activates stimulator of interferon gene (STING)-dependent innate immune pathways and is regulated by mitochondrial membrane potential. J Biol Chem 287, 39776–39788 (2012).

5. Conlon, J. et al. Mouse, but not human STING, binds and signals in response to the vascular disrupting agent 5,6-dimethylxanthenone-4-acetic acid. J Immunol 190, 5216–5225 (2013).

6. Woodward, J.J., Iavarone, A.T. & Portnoy, D.A. c-di-AMP secreted by intracellular Listeria monocytogenes activates a host type I interferon response. Science 328, 1703–1705 (2010).

7. Zhao, B. et al. A conserved PLPLRT/SD motif of STING mediates the recruitment and activation of TBK1. Nature 569, 718–722 (2019).

8. Liu, S. et al. Phosphorylation of innate immune adaptor proteins MAVS, STING, and TRIF induces IRF3 activation. Science 347, aaa2630 (2015).

9. Schlenker, C., Richard, K., Skobelkina, S., Mathena, R.P. & Perkins, D.J. ER-transiting bacterial toxins amplify STING innate immune responses and elicit ER stress. Infect Immun 92, e0030024 (2024).

10. Wu, J. et al. STING-mediated disruption of calcium homeostasis chronically activates ER stress and primes T cell death. J Exp Med 216, 867–883 (2019).

11. Prantner, D., Perkins, D.J. & Vogel, S.N. AMP-activated Kinase (AMPK) Promotes Innate Immunity and Antiviral Defense through Modulation of Stimulator of Interferon Genes (STING) Signaling. J Biol Chem 292, 292–304 (2017).

12. York, A.G. et al. Limiting Cholesterol Biosynthetic Flux Spontaneously Engages Type I IFN Signaling. Cell 163, 1716–1729 (2015).

13. Zhang, X. et al. Sterol Regulatory Element-Binding Proteins and Metabolic Diseases: Mechanisms, Implications, and Therapeutic Strategies. Metab Syndr Relat Disord (2025).

14. Chu, T.T. et al. Tonic prime-boost of STING signalling mediates Niemann-Pick disease type C. Nature 596, 570–575 (2021).

15. Xu, X.J., Reichner, J.S., Mastrofrancesco, B., Henry, W.L., Jr. & Albina, J.E. Prostaglandin E2 suppresses lipopolysaccharide-stimulated IFN-beta production. J Immunol 180, 2125–2131 (2008).

16. Perkins, D.J. et al. Autocrine-paracrine prostaglandin E(2) signaling restricts TLR4 internalization and TRIF signaling. Nat Immunol 19, 1309–1318 (2018).

17. Dennis, E.A. & Norris, P.C. Eicosanoid storm in infection and inflammation. Nat Rev Immunol 15, 511–523 (2015).

18. Woo, S.R. et al. STING-dependent cytosolic DNA sensing mediates innate immune recognition of immunogenic tumors. Immunity 41, 830–842 (2014).

19. Corrales, L. & Gajewski, T.F. Endogenous and pharmacologic targeting of the STING pathway in cancer immunotherapy. Cytokine 77, 245–247 (2016).

20. Corrales, L. et al. Direct Activation of STING in the Tumor Microenvironment Leads to Potent and Systemic Tumor Regression and Immunity. Cell Rep 11, 1018–1030 (2015).

21. Fahey, C.G., Cordova, A.F., Gedeon, P.C. & Barbie, D.A. Targeting STING to generate therapeutic anti-tumor immunity. Cancer Cell (2025).

22. Samson, N. & Ablasser, A. The cGAS-STING pathway and cancer. Nat Cancer 3, 1452–1463 (2022).

23. Hanahan, D. & Weinberg, R.A. Hallmarks of cancer: the next generation. Cell 144, 646–674 (2011).

24. Menendez, J.A. & Lupu, R. Fatty acid synthase and the lipogenic phenotype in cancer pathogenesis. Nat Rev Cancer 7, 763–777 (2007).

25. Currie, E., Schulze, A., Zechner, R., Walther, T.C. & Farese, R.V., Jr. Cellular fatty acid metabolism and cancer. Cell Metab 18, 153–161 (2013).

26. Santos, C.R. & Schulze, A. Lipid metabolism in cancer. FEBS J 279, 2610–2623 (2012).

27. Beloribi-Djefaflia, S., Vasseur, S. & Guillaumond, F. Lipid metabolic reprogramming in cancer cells. Oncogenesis 5, e189 (2016).

28. Al-Khami, A.A. et al. Exogenous lipid uptake induces metabolic and functional reprogramming of tumor-associated myeloid-derived suppressor cells. Oncoimmunology 6, e1344804 (2017).

29. Nielsen, S.F., Nordestgaard, B.G. & Bojesen, S.E. Statin use and reduced cancer-related mortality. N Engl J Med 367, 1792–1802 (2012).

30. Bosch, M., Sweet, M.J., Parton, R.G. & Pol, A. Lipid droplets and the host-pathogen dynamic: FATal attraction? J Cell Biol 220 (2021).

31. Covarrubias, S. et al. High-Throughput CRISPR Screening Identifies Genes Involved in Macrophage Viability and Inflammatory Pathways. Cell Rep 33, 108541 (2020).

32. Pennini, M.E., Perkins, D.J., Salazar, A.M., Lipsky, M. & Vogel, S.N. Complete dependence on IRAK4 kinase activity in TLR2, but not TLR4, signaling pathways underlies decreased cytokine production and increased susceptibility to Streptococcus pneumoniae infection in IRAK4 kinase-inactive mice. J Immunol 190, 307–316 (2013).

33. Sklar, M.D., Tereba, A., Chen, B.D. & Walker, W.S. Transformation of mouse bone marrow cells by transfection with a human oncogene related to c-myc is associated with the endogenous production of macrophage colony stimulating factor 1. J Cell Physiol 125, 403–412 (1985).

34. Perkins, D.J. et al. Salmonella Typhimurium Co-Opts the Host Type I IFN System To Restrict Macrophage Innate Immune Transcriptional Responses Selectively. J Immunol 195, 2461–2471 (2015).

35. Livak, K.J. & Schmittgen, T.D. Analysis of relative gene expression data using real-time quantitative PCR and the 2(-Delta Delta C(T)) Method. Methods 25, 402–408 (2001).

36. Perkins, D.J., Qureshi, N. & Vogel, S.N. A Toll-like receptor-responsive kinase, protein kinase R, is inactivated in endotoxin tolerance through differential K63/K48 ubiquitination. mBio 1 (2010).

37. Li, X., Chen, Y.T., Hu, P. & Huang, W.C. Fatostatin displays high antitumor activity in prostate cancer by blocking SREBP-regulated metabolic pathways and androgen receptor signaling. Mol Cancer Ther 13, 855–866 (2014).

38. Arora, M. et al. Effectiveness and safety of the SREBP1/2 inhibitor, fatostatin, in a preclinical model of metabolic dysfunction-associated steatotic liver disease progression. Eur J Pharmacol 1003, 177890 (2025).

39. Berger, W., De Chandt, M.T. & Cairns, C.B. Zileuton: clinical implications of 5-Lipoxygenase inhibition in severe airway disease. Int J Clin Pract 61, 663–676 (2007).

40. Xu, B., Xing, A. & Li, S. The forgotten type 2 diabetes mellitus medicine: rosiglitazone. Diabetol Int 13, 49–65 (2022).

41. Roberts, Z.J., Ching, L.M. & Vogel, S.N. IFN-beta-dependent inhibition of tumor growth by the vascular disrupting agent 5,6-dimethylxanthenone-4-acetic acid (DMXAA). J Interferon Cytokine Res 28, 133–139 (2008).

42. Zhang, B.C. et al. Cholesterol-binding motifs in STING that control endoplasmic reticulum retention mediate anti-tumoral activity of cholesterol-lowering compounds. Nat Commun 15, 2760 (2024).

43. Wang, Q. et al. The E3 ubiquitin ligase AMFR and INSIG1 bridge the activation of TBK1 kinase by modifying the adaptor STING. Immunity 41, 919–933 (2014).

44. Radmark, O., Werz, O., Steinhilber, D. & Samuelsson, B. 5-Lipoxygenase, a key enzyme for leukotriene biosynthesis in health and disease. Biochim Biophys Acta 1851, 331–339 (2015).

45. Serezani, C.H., Divangahi, M. & Peters-Golden, M. Leukotrienes in Innate Immunity: Still Underappreciated after All These Years? J Immunol 210, 221–227 (2023).

46. Brandt, S.L. et al. Macrophage-derived LTB4 promotes abscess formation and clearance of Staphylococcus aureus skin infection in mice. PLoS Pathog 14, e1007244 (2018).

47. Vila, I.K. et al. STING orchestrates the crosstalk between polyunsaturated fatty acid metabolism and inflammatory responses. Cell Metab 34, 125–139 e128 (2022).

48. Yang, P. et al. Arachidonic acid metabolism in human prostate cancer. Int J Oncol 41, 1495–1503 (2012).

49. Kahnt, A.S., Hafner, A.K. & Steinhilber, D. The role of human 5-Lipoxygenase (5-LO) in carcinogenesis - a question of canonical and non-canonical functions. Oncogene 43, 1319–1327 (2024).

50. Zhou, Q.D. et al. Interferon-mediated reprogramming of membrane cholesterol to evade bacterial toxins. Nat Immunol 21, 746–755 (2020).

51. Brunsting, E.L. & Perkins, D.J. Working in negative space: Type I interferon mediated immuno-modulation through transcriptional suppression in disease and homeostasis. Innate Immun 31, 17534259251367263 (2025).

52. Liston, A. & Masters, S.L. Homeostasis-altering molecular processes as mechanisms of inflammasome activation. Nat Rev Immunol 17, 208–214 (2017).

53. Henriksbo, B.D., Tamrakar, A.K., Phulka, J.S., Barra, N.G. & Schertzer, J.D. Statins activate the NLRP3 inflammasome and impair insulin signaling via p38 and mTOR. Am J Physiol Endocrinol Metab 319, E110–E116 (2020).

54. Pei, G. et al. Cellular stress promotes NOD1/2-dependent inflammation via the endogenous metabolite sphingosine-1-phosphate. EMBO J 40, e106272 (2021).

55. Yu, Y. et al. STING-mediated inflammation in Kupffer cells contributes to progression of nonalcoholic steatohepatitis. J Clin Invest 129, 546–555 (2019).

56. Lv, J. et al. The STING in Non-Alcoholic Fatty Liver Diseases: Potential Therapeutic Targets in Inflammation-Carcinogenesis Pathway. Pharmaceuticals (Basel) 15 (2022).

57. Bai, J. et al. DsbA-L prevents obesity-induced inflammation and insulin resistance by suppressing the mtDNA release-activated cGAS-cGAMP-STING pathway. Proc Natl Acad Sci U S A 114, 12196–12201 (2017).

